# Online learning for orientation estimation during translation in an insect ring attractor network

**DOI:** 10.1101/2021.01.07.425323

**Authors:** Brian S. Robinson, Raphael Norman-Tenazas, Martha Cervantes, Danilo Symonette, Erik C. Johnson, Justin Joyce, Patricia K. Rivlin, Grace Hwang, Kechen Zhang, William Gray-Roncal

**Author notes:** Authors contributed equally (Brian S. Robinson, Raphael Norman-Tenazas).

## Abstract

Insect neural systems are a promising source of inspiration for new algorithms for navigation, especially on low size, weight, and power platforms. There have been unprecedented recent neuroscience breakthroughs with *Drosophila* in behavioral and neural imaging experiments as well as the mapping of detailed connectivity of neural structures. General mechanisms for learning orientation in the central complex (CX) of *Drosophila* have been investigated previously; however, it is unclear how these underlying mechanisms extend to cases where there is translation through an environment (beyond only rotation), which is critical for navigation in robotic systems. Here, we develop a CX neural connectivity-constrained model that performs sensor fusion, as well as unsupervised learning of visual features for path integration; we demonstrate the viability of this circuit for use in robotic systems in simulated and physical environments. Furthermore, we propose a theoretical understanding of how distributed online unsupervised network weight modification can be leveraged for learning in a trajectory through an environment by minimizing of orientation estimation error. Overall, our results here may enable a new class of CX-derived low power robotic navigation algorithms and lead to testable predictions to inform future neuroscience experiments.

**Summary:** An insect neural connectivity-constrained model performs sensor fusion and online learning for orientation estimation.

## MAIN TEXT

## Introduction

A fundamental problem facing both autonomous robots and biological organisms is the task of navigating complex and time-varying environments. In the robotics field, this is a long standing problem that has inspired decades of research into increasingly powerful approaches (*1*). The navigation challenge can be subdivided into components such as mapping (*2, 3*), path planning (*4*), sensing, and actuation. Critical components include state estimation of the robot using sensor data and sensor fusion, including classic approaches such as Kalman filtering, along with its nonlinear extensions (*5, 6*) as well as loop closure, where storage of processed sensory data is used to recognize the return to specific locations in an environment (*2, 3*).

Real time state estimation for low-power platforms, such as small drones, remains dominated by visual odometry (*7*). Many visual odometry approaches for state estimation exist (*8*), including high performing Visual-Inertial Odometry (VIO) systems for state estimation on rapidly moving platforms (*7*). In general, these approaches require simplifying assumptions and linearization of the system dynamics, as a full solution of the state estimation problem for noisy data requires propagation of the full conditional probability distribution (*9*), though this can be addressed with more computationally intensive approaches such as particle filtering (*10*). Nonlinear approaches which adapt to changes in the environment remain an ongoing research challenge. Additionally, visual odometry approaches do not perform loop closure and are prone to error accumulation when used in isolation. In particular, observed visual features that have been previously encountered are not used to update pose estimation.

Algorithms for visual approaches to Simultaneous Localization and Mapping (SLAM), which perform loop closure, are extensively used on higher power platforms (*11*–*14*). Real time, efficient implementation of visual SLAM systems on embedded mobile platforms remains a major engineering challenge (*15, 16*), often requiring custom hardware designs for bundle adjustment and loop closure (*17*). There are also emerging visual navigation and visual SLAM approaches based on deep learning, including self-supervised approaches (*18*). This has resulted in powerful approaches for monocular visual odometry (*19*), and also novel strategies such as vector navigation on representations with grid-cell like responses (*20*) and predictive navigation in complex environments (*21*). In general, however, training of the neural networks utilized in visual navigation are not performed during robotic system deployment due to computational requirements for training and the number of required training samples.

Neuromorphic processing is a promising path forward for embedded neural network navigation approaches considering the ability for on-chip training as well as the incorporation of low-latency event-driven sensors. Event-driven cameras and processing have been shown to produce low-latency, high-performance VIO systems for quadcopters (*22*). This path is even more promising due to the development of general purpose neuromorphic hardware, such as the spinnaker (*23*), IBM’s TrueNorth chip (*24*), and the Intel Loihi (*25*). The Loihi is particularly promising for robotics applications, as it allows on-chip learning and has demonstrated efficiency at the classification of dynamic data in applications such as keyword spotting (*26*) and classifying olfactory signals (*27*). Navigation strategies utilizing event-driven data and neuromorphic processing are particularly compelling for the potential to provide low-latency, adaptive navigation solutions with efficient implementations and motivate further algorithm development.

Biological systems are a natural source of inspiration for navigation algorithm development for neuromorphic processing, considering their navigation capabilities and low power utilization. Mammalian systems are among the most studied in neuroscience in relation to navigation with research breakthroughs over the past decades identifying several specialized cell types including head direction cells (*28*), grid cells (*29*), and place cells (*30*). There have been several approaches utilizing these representations and additional specialized mammalian cell types (*31, 32*) to propose algorithms for neuromorphic robotic navigation (*33*–*38*). All of these neurons represent space in an allocentric coordinate system, which is in reference to a world frame, rather than in an egocentric coordinate system, centered to the animal’s body or the sensors as commonly seen in low-level sensory processing. Additionally, these neurons generally derive their activities by combining two types of inputs, one is based on integration of self-motion information, and the other is based on sensory signals from the environment.

Head-direction cells are found across different mammalian species (*31*) such as rat, mouse, chinchilla (*39*), macaque monkey (*40*), and flying bat (*41*) as well as non-mammalian species such as the fly, *Drosophila melanogaster* (*42*). The head-direction cells are conceptually the simplest among the spatial neurons because the internal representation is only one-dimensional with ring topology. Currently the leading theoretical models for the head-direction cells are ring attractor networks (*43*–*45*). Although many models have been proposed (*46*–*49*), they all share the same basic principle of using a stable equilibrium state of a ring network, which we call an activity bump, to represent the current internal sense of direction. The input from angular velocity of head movement serves to drive the activity bump around the ring of cells, and the input from sensed landmarks serves to anchor the peak position of the bump to the learned position. Although the mammalian head-direction systems have been intensely studied in the ensuing decades since its initial discovery, connections from the visual or vestibular systems to the conceptual ring topology have to go through multiple bilateral structures, including dorsal tegmentum, lateral mammillary body, anterior thalamus, and postsubiculum (*50, 51*); consequently the detailed connectome has never been established. Thus these theorized networks are a vast simplification, not constrained by experimental observations or known anatomical connectivity. Existing robotic studies inspired by the mammalian head direction systems are conceptual due to the unknown mechanisms for integrating angular velocity and visual feature cues at an observed cell-type level (*35, 52*–*57*). Additionally, the presence of close landmarks, which have sensed visual features that vary based on position (as opposed to landmarks at an infinite distance on the horizon), have been less well studied in mammalian head direction computational models (*58*) and represent a necessary challenge in creating an orientation estimation system in cluttered environments during translation.

Compared to mammalian systems, insect systems are a compelling source for navigation algorithm development because of the low size, weight and power (SWaP) of insects, the relative simplicity of the networks, and an enhanced mechanistic understanding, whereby the detailed connectivity patterns are known. Recent experimental breakthroughs in *Drosophila* have identified a candidate set of “compass” neurons involved in a ring attractor network for representing heading direction in a neural region called the central complex (CX). Elucidating the structure and function of *Drosophila* is an active area of study and large-scale wiring diagrams are being reconstructed, providing a rich substrate for analysis, model refinement and validation (*59, 60*). Compared to the theoretical plausibility of a ring attractor for heading direction in mammalian systems, experiments have observed and quantified an activity bump corresponding to the insect’s heading direction (*42, 61*). Furthermore, neurons responsible for providing input into the ring attractor for angular velocity(*62, 63*) and visual features (*64, 65*) have been characterized experimentally. Correspondingly, the detailed anatomical connectivity patterns between neurons involved in the network have been observed (*66*) and investigated for their computational role (*67*– *69*) at the level of individual neuron type to support ring attractor dynamics. A unique opportunity exists, therefore, to derive mechanistic inspiration for a navigation system utilizing detailed connectivity patterns observed in *Drosophila* for representing heading direction in a ring attractor network.

In order to utilize visual landmark cues from an environment for orientation estimation, a mapping must be learned between the location of visual features on a sensor to an orientation referenced to a world frame coordinate system. Theoretical models have been proposed for how Hebbian unsupervised learning can lead to the strengthening of connections between neurons receptive to visual features and heading direction ring attractor neurons that are co-active in order to learn the mapping between landmark cues and an orientation estimate (*43, 70*). Hebbian learning is a general type of synaptic mechanisms of associative learning well studied in neuroscience and recent experimental evidence in *Drosophila* support a role for Hebbian learning for the mapping between neurons representing visual features and the compass neurons (*65, 71*).

While there has been compelling insight into the *Drosophila* heading direction system at a cell-type level, an outstanding question is how amenable this approach may be for robotic applications that include a trajectory through an environment. In particular, the performance of the heading direction system has not been quantified when using cell-type specific connectivity patterns. Additionally, due to constraints of experimental approaches, both empirical observations and existing computational models have been limited to single location trajectories where there is rotation without translation. In particular, the coordinated learning of a visual feature mapping while traversing a trajectory through an environment is a necessary and challenging function for such a network that is not currently understood.

The key contributions of this work are to develop and evaluate a model for how a *Drosophila* connectivity-constrained network can perform both sensor fusion and online learning for estimating orientation in a trajectory through an environment. We evaluate the model through trajectories with noisy inputs in a simulated environment and investigate performance with measurements from a robotic platform. Furthermore, we propose a theoretical understanding of how distributed online unsupervised learning can be leveraged for learning in a trajectory through an environment in coordination with a ring attractor network.

## Results

### Connectivity-constrained network for sensor fusion and online learning

Our model transforms angular velocity and visual features into a fused representation of orientation utilizing five populations of neurons as specified in Fig 1A, which are based on connectivity patterns of neuron types observed in *Drosophila*, and with plastic synaptic connections that enable the learning of visual landmarks. The five types of modeled neurons all synapse in either the protocebral bridge (PB) or ellipsoid body (EB) regions of the CX and include: 1) Ring neurons which are receptive to visual inputs, 2) PB-EB-Noduli (P-EN) neurons which receive angular velocity inputs, 3) EB-PB-Gall neurons (E-PG) neurons, 4) PB-EB-Gall (P-EG) neurons, and 5) Intrinisic neurons of the PB (Pintr), also referred to as Δ7 neurons. Orientation is encoded in a population of E-PG neurons or “compass” neurons, which have been observed in experimental studies to have maximal activation in a position along the anatomical circumference of the ellipsoid body which rotates corresponding to the orientation of the fly. The modeled population of 18 E-PG neurons is divided into left and right hemispheric groupings depending on whether the neuron projects to the left or right hemisphere of the PB. We utilize an orientation representation scheme where each of the 9 E-PG neurons per hemisphere is maximally active at a distributed preferred angle as demonstrated in Fig 1C.

**Fig 1.**
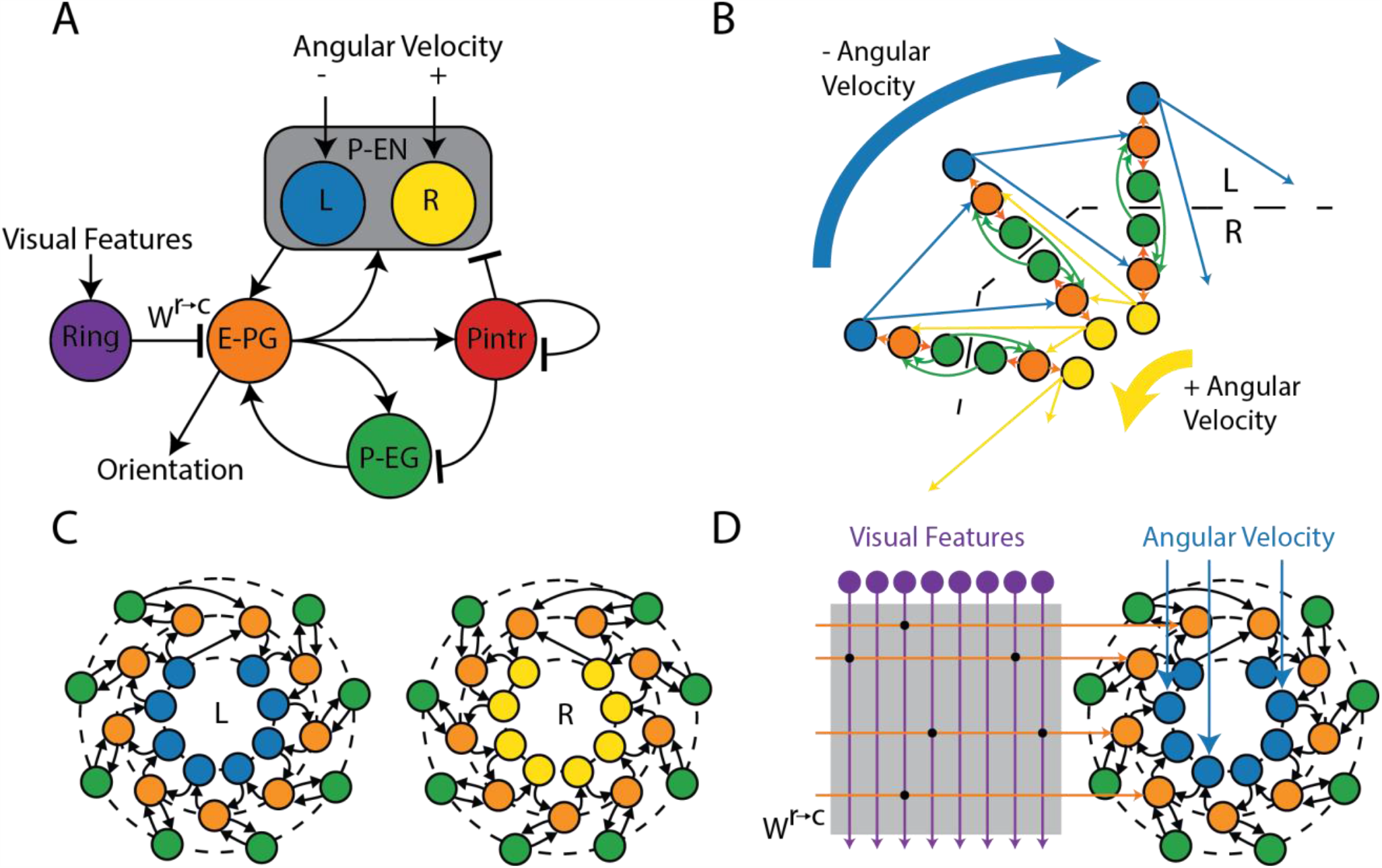
*Drosophila* CX ring attractor model for sensor fusion and environmental learning. (**A**) Five neuron types found in the *Drosophila* central complex (CX) included in the ring attractor model. Input is provided to the model as visual features to the ring neurons and angular velocity to the P-EN neurons. P-EN neurons are preferentially activated by positive and negative angular velocities according to if they are located in the left or right hemisphere of the protocerebral bridge. Orientation is encoded in E-PG compass neurons. All neuron types are excitatory except for Pintr and ring neurons which provide inhibition. (**B**) The primary repeated pattern in the connectivity constrained ring attractor network organized by E-PG neuron projecting to glomeruli in the left and right hemisphere of the protocerebral bridge. Local recurrent excitation enables the sustained activation of an activity bump, with positive or negative angular velocities shifting the center of the activity bump across the ring attractor network. (**C**) The connectivity pattern utilized for all excitatory modeled neurons separated by protocerebral bridge hemisphere. Note that there is one additional modeled E-PG neuron per hemisphere than other neuron types. (**D**) A weight matrix, ***W***^***r***→ ***c***^, is learned, which specifies the strength of connections between ring neurons and compass neurons.

A stable bump of activity in E-PG neurons can persist given the recurrent excitation to keep the bump active along with inhibition provided by PIntr neurons to prevent runaway excitation. Local recurrent excitation for maintaining bump activation in E-PG neurons is mediated by both P-EN and P-EG neurons. Positive angular velocity is encoded in the P-EN neurons in the right hemisphere, which when active, shifts the activity bump in the counter-clockwise direction. Conversely, negative angular velocity is encoded by P-EN neurons in the left hemisphere which shifts the activity bump in the clockwise direction (Fig. 1B).

Sensed visual features represented in ring neuron activation patterns can update the activity bump in E-PG neurons according to their connectivity weights (Fig. 1D). The intuition behind the online learning of visual features is that the activity bump is initially driven by angular velocity signals which serves as a noisy teaching signal to update a matrix of weights, *W*^*r*→ *c*^, between ring neurons and “compass” E-PG neurons. Each individual element, *w*_*nm*_, is the weight between the m^th^ ring neuron and the n^th^ E-PG neuron. The mechanism utilized for learning is a Hebbian learning rule, where the co-activation of neurons increases the effective connection weights between ring and E-PG neurons. In particular, a presynaptically-gated Hebbian learning rule is used to modify weights (*71*),

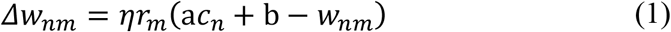

where *r*_*m*_ is the activation of the m^th^ ring neuron, and *c*_*n*_ is the activation trace of the n^th^ E-PG neuron. The rule is parameterized with a learning rate, *η*, as well as additional constants a and b. All utilized *w*_*nm*_ weights are assumed to be negative, as in previous computational studies (*67*) given the inhibitory characteristics of ring neurons (*65, 72*), so the maximum effective weight is zero, which can activate E-PG neurons through disinhibition. The model’s orientation estimate is calculated as the center of the activity bump utilizing the preferred direction of each E-PG neuron to create a linear decoder applied to filtered spiking events. Additional details for model implementation can be found in the methods section.

### Model performance with rotation

An initial evaluation of the network is performed in the case of a simple rotation in an environment without translation (Fig 2A). Ring neurons encode visual input, with a separate set of ring neurons responsive to each landmark and each individual ring neuron with a receptive field tiled across a 270° field of view. As the simulated agent and field of view rotates, the index of the most active ring neuron in each sub-population shifts (Fig 2B). A corresponding shift in a bump of activity is observed in E-PG, P-EN, and P-EG neurons (Fig. 2C-E), which appears as two bumps given the separate indexing of left and right hemisphere neurons. Angular velocity input is provided to the model by injecting current in the P-EN neurons, where a counter-clockwise rotation in the simulation corresponds to an observed increase in activation of right P-EN neurons (Fig. 2C). Throughout the trial, there is consistent activation of Pintr neurons (Fig. 2D), which provide inhibition to the network and prevents the model from going into a state of runaway excitation. The estimated orientation is decoded during the course of the trial from E-PG neurons as the center of the bump of activation in relation to the preferred direction of each E-PG neuron.

**Fig 2.**
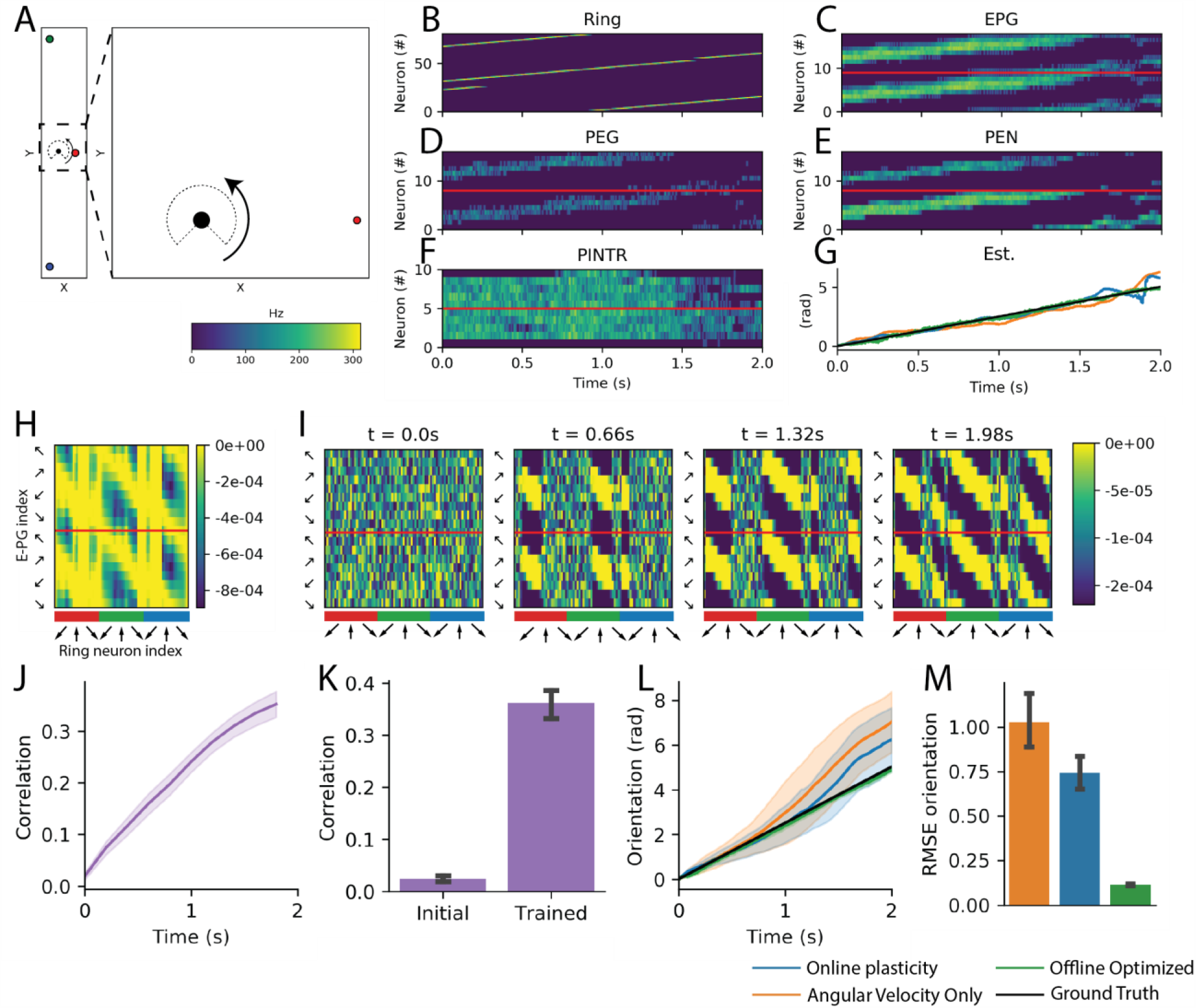
Sensor fusion and online learning with rotation in simulation. (**A**) Visual and angular velocity inputs to network correspond to a rotation in a simulated environment with landmarks and a 270° field of view. (**B-F**) Neuron activation over time in each population supports sensor fusion of inputs into an activity bump that shifts in the E-PG neuron population during rotation. **(G)** Orientation estimation decoded from E-PG neuron activity (best seed example) with three evaluated model configurations tracks the ground truth orientation. (**H**) Offline optimized ***W***^***r***→ ***c***^ specifies an optimal mapping from visual landmarks encoded in ring neurons (x axis) to E-PG “compass” neuron activation (y axis). Ring neuron indices are arranged sequentially according to receptive field center for each colored landmark. (**I**) In the model configuration with online plasticity, weights evolve over time to a similar structure as offline optimized weights. (**J-K**) Correlation between online learned weights and the offline optimized solution increases over time. (**L**) Estimated orientation over time drifts less with online plasticity then with angular velocity alone. (**M**) Average orientation RMSE. All group level analyses in (**J-M**) are performed for 50 simulated seeds. (**J, K, M**) have 95% bootstrapped confidence bounds, (**L**) has standard deviation confidence bounds.

Three model configurations are compared in order to evaluate sensor fusion and online learning 1) initialization of *W*^*r*→ *c*^ weights to an optimal set of offline calculated weights, 2) online learning of modeled weights according to Eq. 1 after random initialization, and 3) setting all *W*^*r*→ *c*^ weights to zero such that angular velocity is the only input into the model. Due to the stochasticity of the neuron model, a total of 50 simulation seeds are used to evaluate performance. An example of the decoded orientation from all three model configurations is shown in Fig. 2G, where all of the model configurations are able to track the orientation over time (with the optimal offline weights calculated weights having the best performance). Given the evaluated trajectory where there is no translation, any pair of landmarks that are offset at an angle wider than the gap in the visual field would provide enough visual input to perform orientation estimation. The optimized weights (Fig. 2H) are calculated with regularization which effectively select a subset of model ring neurons to utilize for angle estimation. The weights that are learned over the course of the trial (Fig. 2I), have a similar banded structure to the optimized weights and increase in correlation over the course of the trial to an average value of 0.36 (Fig. 2J). Factors limiting higher correlation include 1) the effective “teaching” signal in the online training is from integrating angular velocity cues which are inherently noisy as well as 2) the inherent differences between the training approaches. Estimated orientation error accumulates over the course of the trial in the angular velocity only case (Fig. 2L), which is reduced when there is online learning. Overall, the average orientation root mean squared error (RMSE) is significantly reduced by 28% (1.03 to 0.74 radians, P=.0023, Mann-Whitney *U*-test) with the online learning verses angular velocity alone, compared to an 89% reduction with the set of optimized weights. Sources of error in the weight optimization process include: 1) *W*^*r*→ *c*^ is optimized with regards to the feedforward visual input alone and target E-PG activity which does not explicitly compensate for the recurrent activation from P-EG and P-EN neurons, 2) there is an inherent time lag in spiking neuron activation given the 20ms modeled time constant with the leaky integrate and fire neurons, 3) the orientation is decoded from an estimate of the center of the bump from spiking activity of 18 neurons. Nevertheless, a RMSE of 0.11 radians (6.3°) in the optimized weights demonstrates that this network with a limited number of spiking neurons and recurrent excitation can effectively perform sensor fusion for accurate orientation estimation.

### Model performance with translation

The network is next evaluated in a more challenging trajectory that involves translation through the simulated environment, where there is no longer an invariant transformation between egocentric visual landmark features referenced to a sensor frame and an orientation referenced to an allocentric world frame coordinate system (Fig. 3). The bump of E-PG activation follows the time-varying orientation in the trajectory and is able to be decoded into an accurate orientation estimation over time (Fig. 3A-B). In the trajectory through the environment, there are two distant landmarks, which individually have a relatively invariant transformation between egocentric and allocentric representations, along with a proximal landmark which has a time-varying transformation between egocentric and allocentric representations (Fig. 3C).

**Fig 3.**
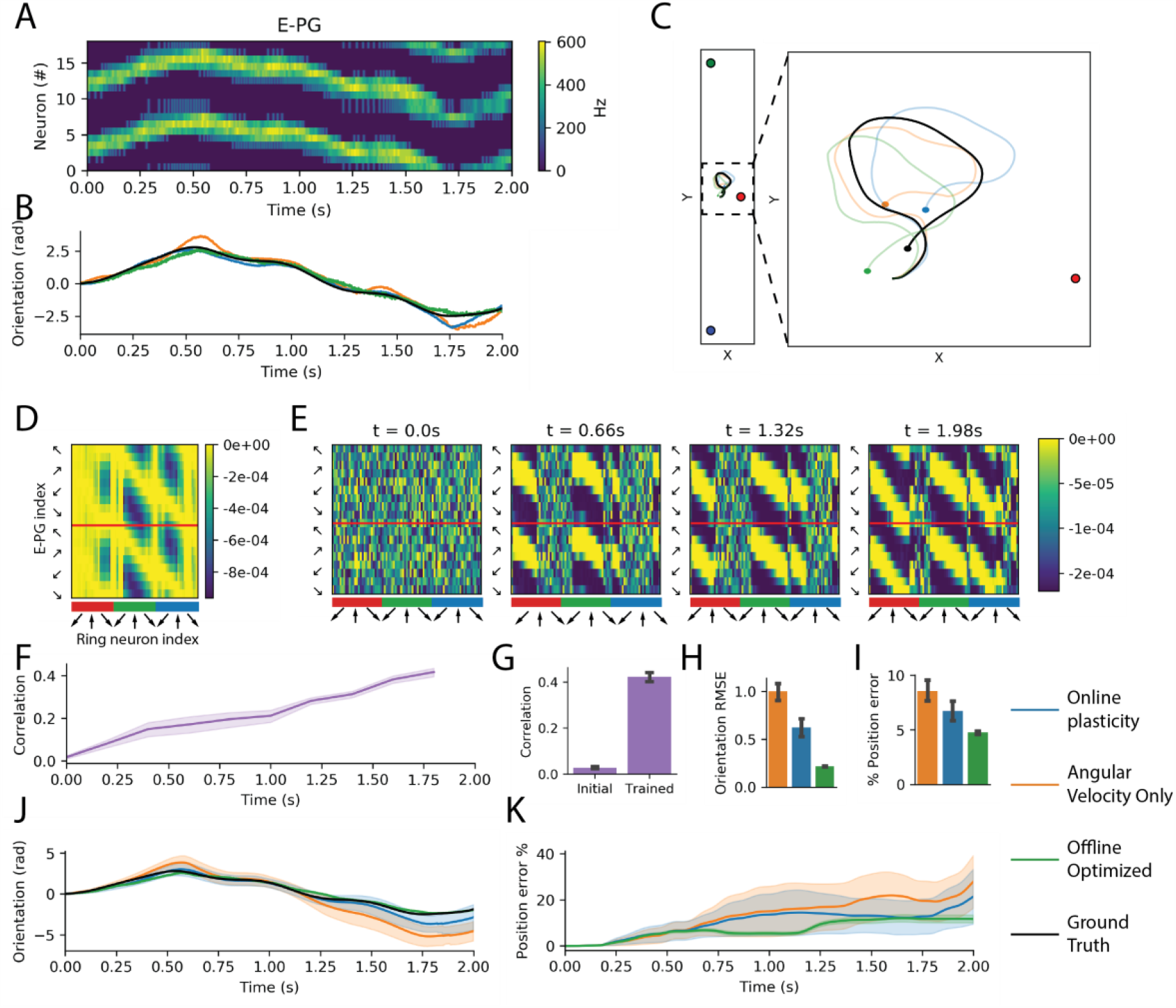
Sensor fusion and online learning with translation in simulation. (**A**) Shifting of the activity bump in E-PG neurons over time in the model configuration with online learning corresponds to changes in orientation. (**B**) The decoded orientation estimation (best seed) with three evaluated model configurations corresponds to the ground truth trajectory. (**C**) In the simulation, there are two distant landmarks and one close landmark which provide visual inputs to the network. Position estimates over the course of a trajectory with path integration approximate ground truth. Final positions in the trajectory estimates are denoted with a circle. (**D**) Offline optimized ***W***^***r***→ ***c***^ has features preferentially tuned to distant landmarks. (**E**) Evolution of weights during online plasticity share overall structure with offline optimized weights. (**F-G**) Correlation between online learned weights and the offline optimized solution increases over time. (**H**) Average orientation RMSE with online plasticity is between offline optimized weights and angular velocity alone model configurations. (**I**) Improvement in position error (average position error as a percentage of path length) additionally observed with online plasticity versus angular velocity alone. In orientation estimates over time (**J**) and position error as a fraction of path length over time (**K**), error accumulates less with online plasticity than angular velocity alone. All group level analyses in (**F-K**) are performed for 50 simulated seeds. (**F-I**) have 95% bootstrapped confidence intervals, (**J-K**) have standard deviation confidence intervals.

The set of offline optimized weights for the network (Fig. 3D) preferentially selects visual features corresponding to the distant landmarks. The evaluated network configuration with online plasticity learns weights with increasing correlation over time to the optimal weights (Fig. 3E-G) and reaches a maximum correlation of 0.42. When compared on multiple simulated seeds, similar general trends for orientation estimation accuracy are observed between network configurations as to the rotation only trajectories (Fig 3. H, J), where the orientation RMSE for online learning is between the angular velocity only case and the optimal set of weights. The RMSE of the angular velocity only network configuration in this trajectory with translation is similar to the simple rotation case (1.00 vs. 1.03 radians), but is compensated further by online learning (38% vs. 28% reduction in error).

Position estimation with path integration is more challenging than orientation estimation because of the accumulation of error driven by orientation estimation error. Similar trends are observed with average position estimation error (as a percentage of path length), where online learning has a position estimation error between the angular velocity alone and the optimized weights network configuration (6.7% vs. 8.6% and 4.8%, Fig. 3I). The decrease in average position error with online learning vs. angular velocity alone is 22% (P= 0.0021, Mann Whitney U-test). Sources of error for position estimation for path integration include the aforementioned error accumulation from orientation estimation that could be further reduced with neuro-inspired (*73*) or traditional approaches (*74*).

Overall, network simulation results with translation demonstrate that an accurate allocentric estimation of orientation referenced to a world frame can be estimated with egocentric visual features despite the challenges of transforming information over time from proximal landmarks. The model configuration with online learning is correlated with the optimized set of weights and improves orientation and position estimation by an average of 38% and 22% respectively versus a configuration with angular integration alone. Simulation results suggest that improvements to the online learning rule (Eq. 1) may be beneficial to effectively select visual features from distal landmarks to increase accuracy further.

### Model performance on a robotic platform

The model for orientation estimation and position estimation is extended from simulated environments to measurements from a wheeled robotic platform (Fig 4A) in an arena with colored landmarks (Fig 4B). Visual inputs are measured from a camera with two separated sensors offset at 180°, where the relative angle offset of landmarks is calculated from blob detection on color-masked images (Fig 4C). The relative angle offset detected for the center of each of the green, yellow, and red colored landmarks are used to drive the activation of ring neurons (Fig 4D) according to their receptive fields, which are mapped equivalently to the three populations of ring neurons mapped to simulated landmarks across a 270° field of view. Given the less than 360° field of view on the camera sensors, there are ring neurons selective to outside the camera’s field of view which will never activate. While several different neural visual encoding schemes are possible which would likely improve performance, mirroring the simplified visual encoding from simulations enables straightforward comparison between simulated and physically measured model performance. An added source of noise in the visual feature measurements is the intermittent dropping of recorded image frames due to maximum disk-writing speeds, which is apparent in the lack of ring neuron activity at intermittent periods, which further tests the network’s performance in relation to sensor fusion. The angular velocity measurements to drive the activation of P-EN neurons (Fig 4E) are derived from an on-board IMU sensor. Note how the activation is greater on the right P-EN neurons given the positive angular velocity measurements. The right P-EN neurons are still active, however, because of recurrent excitation in the network.

**Fig 4.**
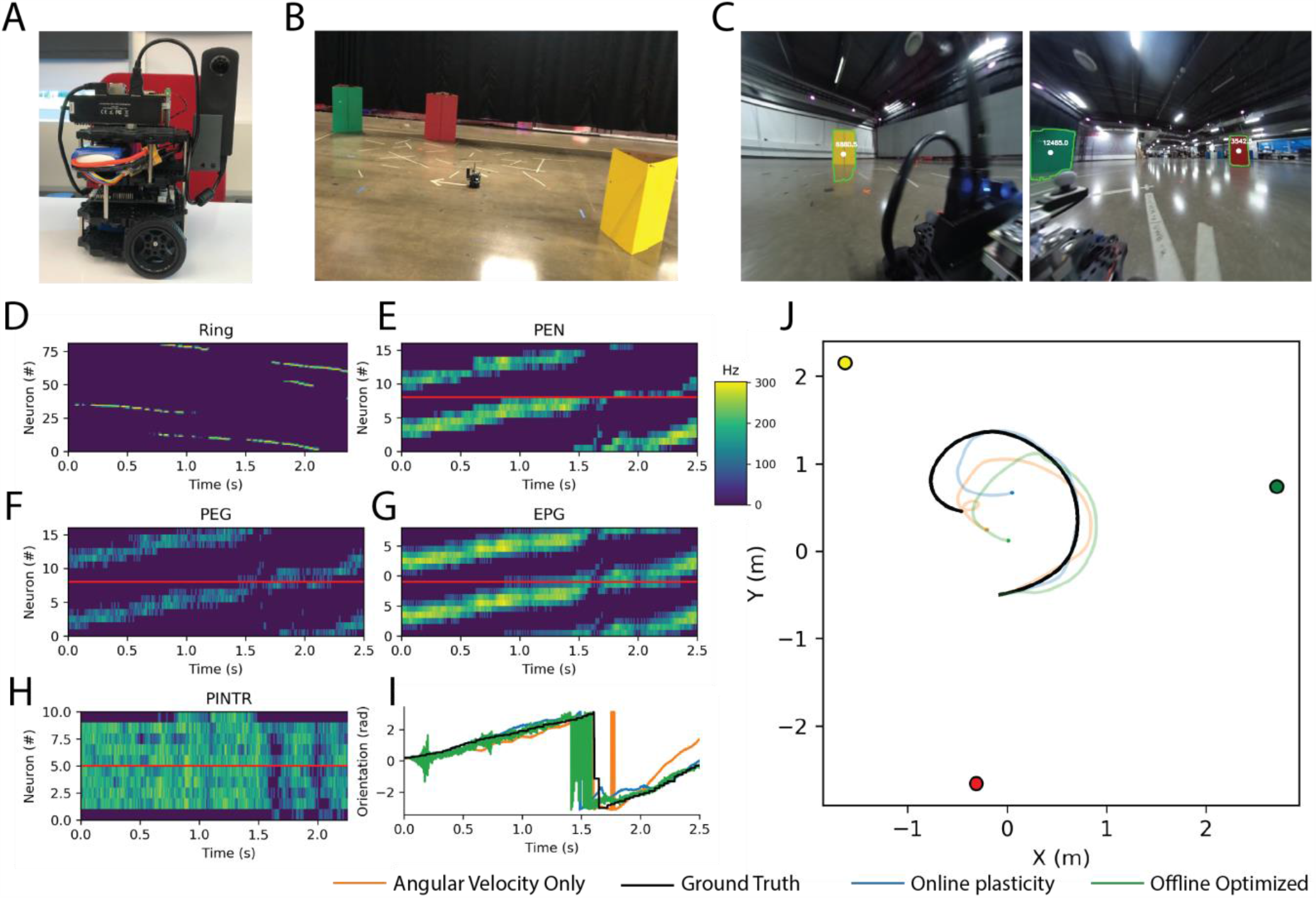
Model utilized with a robotic platform. (**A**) Modified Robotis Turtlebot3 “Burger,” equipped with a Nvidia Jetson TX2, and a Ricoh Theta S 360° camera. (**B**) Physical arena with colored landmark cues. (**C**) Example camera measurements with blob detection utilized to drive activation of visual neurons. (**D-H**) Neuron activation over time in each population driven by processed sensory data from the robotic platform, where visual landmark features encoded in ring neurons (**D**) drive shift in activity bump in E-PG neurons (**G**) in model configuration with online plasticity. (**I**) The estimated orientation from each model configuration (best seed) tracks the ground truth orientation with errors in the offline optimized weights at periods of prolonged visual dropout. (**J**) In the arena, the ground truth trajectory is between all of the landmarks with position estimates from each model configuration that share features of the overall trajectory (best seed). Final positions in the trajectory estimates are denoted with a circle.

The center of the bump of activity in the P-EG and E-PG neurons shifts over the time (Fig 4F-G), which leads to a decoded orientation that follows the ground truth orientation (Fig 4I). A position estimate is performed with path integration, where a ground truth measure of linear velocity is integrated with the orientation estimate (Fig 4J). Similar to the model evaluation in simulation, a comparison of the model performance with several seeds of neural simulations with three model configurations is performed with the plasticity model, the angular velocity only model, and the optimized weights. An example of the comparison between the optimized weights and the online learning of weights is shown in Fig 5A-B, which again share a banded structure. The correlation of the optimized and online learned weights over time is quantified over 50 seeds in Fig 5C-D, where the correlation monotonically increases over time to a maximum average value of 0.29. The orientation error accumulates over time fastest in the angular velocity only case with a slight decrease in error accumulation with online learning (Fig. 5G). Overall, the orientation RMSE is 14% less with online learning vs. angular velocity alone (average 0.80 vs. 0.93 radians, P=0.014, Mann-Whitney *U*-test), however the optimized weights have the lowest error. The position similarly accumulates over time in all comparisons, with a notable increase in position error at 1.5s in the weight optimized case due to a prolonged period of dropout of visual features (Fig. 5H). A minor decrease in the average position error with the online learning versus the angular velocity case alone is observed.

**Fig 5.**
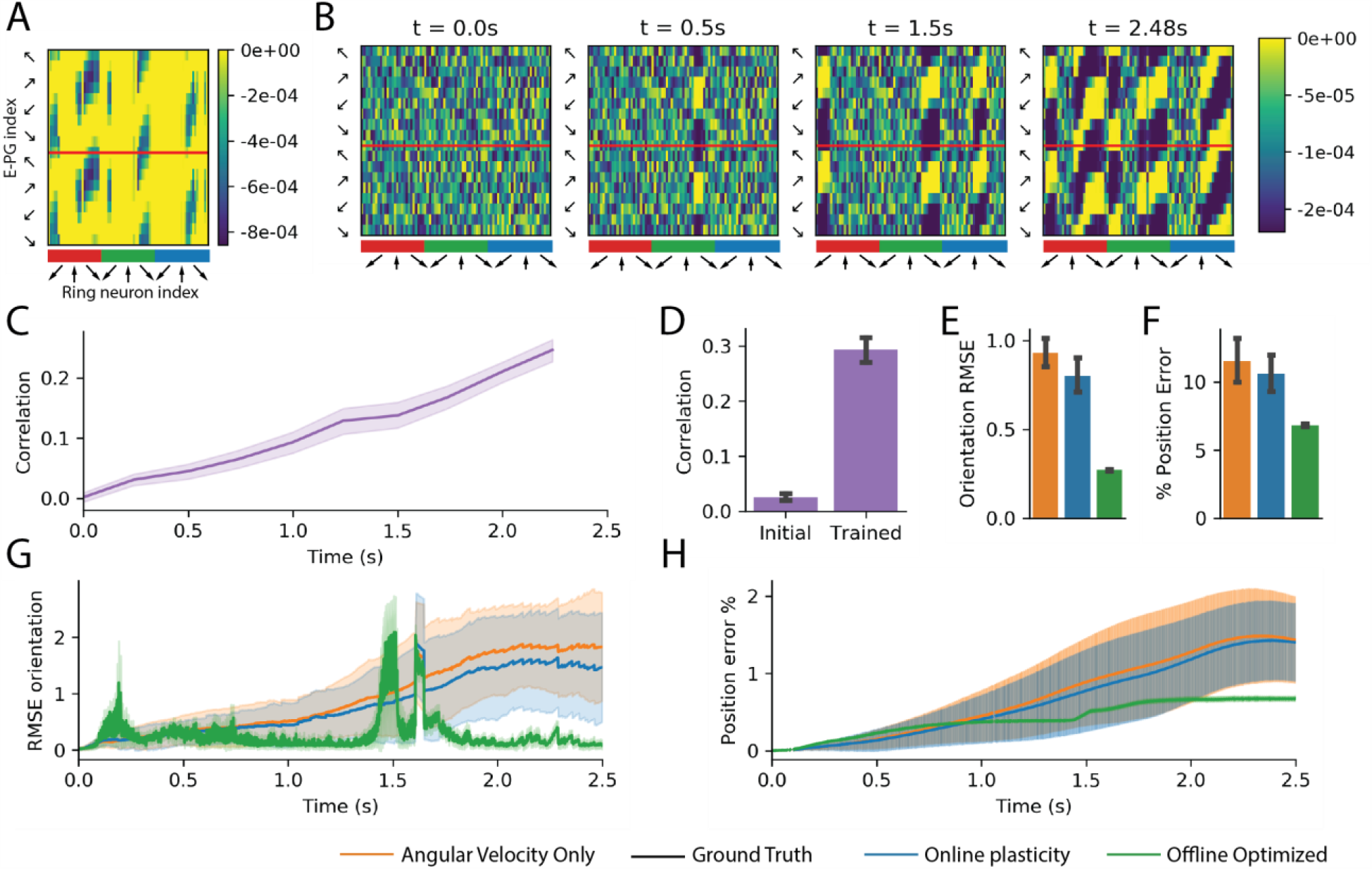
Model evaluation on a robotic platform. (**A**) Offline optimized ***W***^***r***→ ***c***^ has a banded structure which represents an optimal set of weight to map visual features in ring neurons to orientation representation in E-PG neurons. (**B**) Evolution of weights during online plasticity increases in similarity to offline optimized weights over time. (**C-D**) The correlation between online learned weights and the offline optimized solution increases over time. (**E**) The average orientation RMSE (in radians) is less with online plasticity than angular velocity alone. (**F**) Average position error as a percentage of path length has similar trends as orientation estimation with less differentiation between online learning and angular velocity alone. (**G**) In estimated orientation RMSE over time, error increases in periods with visual feature dropout with offline optimized weights. (**H**) Position error as a percentage of path length over time. All group level analyses in (**C-H**) are performed for 50 simulated seeds. (**C-F**) have 95% bootstrapped confidence intervals, (**G-H**) have standard deviation confidence intervals.

Opportunities to improve position error estimation include: 1) improvements to decrease the angular velocity alone orientation estimate that is effectively the self-supervised teaching signal of the network, 2) improvements to the online learning rule, 3) longer periods of exploration in the environment. Overall, these results a critical proof of principle, demonstrating the viability of an insect connectivity-constrained orientation estimation model with learning applied to a robotic platform and identify areas of continued research in order to be able to leverage an insect-inspired solution on a robotic platform.

### Online learning analysis

In all experiments, weights derived from online learning are correlated with an optimal solution and improve orientation accuracy after a single trajectory. A pressing question for robotic translation is how to enhance estimation accuracy to further minimize error accumulation.

One approach is to identify the objective function that the utilized learning rule is minimizing. Upon further inspection (see Methods Eq. 5-6), it follows that Eq. 1 is minimizing the overall objective function,

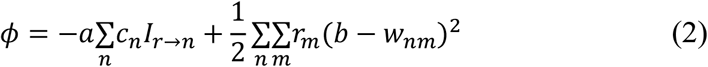

where *I*_*r*→ *n*_ = ∑_*m*_ *r*_*m*_ *w*_*n,m*_. The first term, 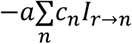, minimizes the objective function when the E-PG neuron activity and the total input current from all of the ring neurons for each compass neuron are aligned. The second term, 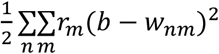, effectively acts as a regularization term on the weight values, which is minimized the closer each weight value is to *b*. While the learning rule (Eq. 1) effectively maximizes the overlap from the input current from ring neurons and the compass neuron activation, it is not directly optimizing an objective function to minimize the squared error of the orientation estimate. An example of this is how the online learning model (Eq. 1) did not preferentially select distal over proximal landmark features. The success of utilizing a scaled version of the offline optimized weights solution in preferentially selecting distal landmark visual features in simulation (and increasing accuracy overall in simulations and on the robotic platform), lends support to using a set of directly optimized weights that minimize squared orientation error in future approaches. In order to directly minimize the objective function of the scaled squared orientation estimation difference, as in the offline optimized weight calculation, we can define the objective function for each compass neuron,

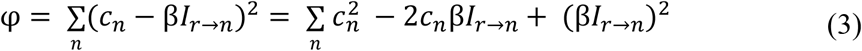

with a scaling factor, β. Upon further inspection, it follows that in order to minimize the objective function, φ, (see Methods Eq. 7-8), a weight update rule is

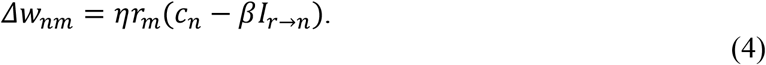

This rule has a commonality to the previously utilized rule (Eq. 1), where weight modifications are presynaptically gated; however, there is an additional term, *I*_*r*→ *n*_, that is utilized. Note how in the expanded objective function term in Eq. 3, there is a similarity between terms *c*_*n*_β*I*_*r*→ *n*_with the first term in the pre-synaptically gated objective function (Eq. 2) to maximize the overlap between the *c*_*n*_ activity and the current from the ring neurons, however in Eq. 3, there are additional terms that penalize the individual compass neuron activation and input current levels. We propose the learning rule above (Eq. 4) for utilization in future online robotic applications to enable an improvement in accuracy in learning an environmental mapping. Each of the terms utilized in Eq. 4 is biologically plausible, which could motivate future experimental evaluation. Furthermore, each term is a local variable specific to pairs of synaptically coupled neurons, which would be amenable for implementation in distributed learning on neuromorphic hardware.

## Discussion

Our motivation is to investigate the potential for leverage details from *Drosophila* neurobiology and neuroanatomy for sensor fusion and online learning for orientation estimation as a basis for future low SWaP neuromorphic robotic navigation approaches. We also hope to better understand the underlying neural circuits toward additional model development and exploration. We develop a model for *Drosophila* sensor fusion and online learning in a cell-type connectivity-constrained model for orientation estimation and environmental learning that integrates angular velocity measurements and visual features. Through a series of experiments in simulated environments, we demonstrate that improvement in orientation and position accuracy estimation is possible with online learning of visual features (versus angular velocity alone) over a single trajectory and that online learned weights are correlated with a set of offline calculated optimal weights. The network model is adapted for use with sensors from a robotic platform and is demonstrated to have similar trends in performance with increased accuracy with online learning over a single trajectory. Finally, a theoretical understanding and novel weight update rule for distributed online learning with local variables is proposed that can be utilized to minimize estimation error.

There are many neuroscience-inspired robotic approaches for navigation that are largely inspired by the cell types observed in mammalian systems, e.g. (*33*), with a subset of these utilizing a ring attractor to represent heading direction, e.g. (*54*). Compared to mammalian-inspired approaches for navigation, insect-inspired approaches are promising for translation to low SWaP robotic platforms considering the relative compactness and simplicity of the underlying neural circuits, the enhanced details observed regarding the cell-type specific connectivity patterns, and the rich set of experimental evidence demonstrating the dynamics and online learning at the scale of an entire neural region. Our results are the first to demonstrate how an insect cell-type connectivity-constrained model for ring attractor heading direction estimation can be operationalized on a robotic platform.

Outside of robotic investigation, there has been a wider array of modeling results investigating the computational principles of mammalian and insect navigation, with a subset of these focusing on heading direction estimation with a ring attractor network (*43, 70*). An outstanding challenge to these approaches investigated in this work for heading direction estimation is how to learn and integrate visual information from cluttered environments, where the relationship between visual features and heading direction changes through a trajectory in an environment (*58*). The network model utilized here is a novel *Drosophila* cell-type connectivity-constrained model for sensor fusion with online learning evaluated in the context of translation through an environment with local visual features.

Our results underscore the importance of translational movements in the representation of head direction. A hallmark of a typical mammalian head-direction cell is that the preferred direction of the cell is the same regardless of the animal’s location. In the *Drosophila* heading system, experiments have been carried out in a position-fixed setup with virtual reality visual display that updates with rotation but does not simulate translation through an environment. It is yet to be experimentally verified whether the preferred direction is kept the same in different locations. Prior computational modeling studies of the system were also based on restricting the head to a single fixed location. Our results show that once the spatial location is allowed to vary, keeping the preferred direction consistent across spatial locations becomes a more difficult learning problem. We also found that landmarks at a distance help anchor the heading system better than nearby landmarks through analysis of optimized visual feature weights, which is intuitive because a distant landmark provides a more consistent signal at different positions in an environment. By contrast, when the head is fixed to a single location, the angular movements of landmarks are identical regardless of their distances. Our results demonstrate that in the insect heading direction research, it is important to test the system with multiple locations, both in real and virtual environments.

Our results include an investigation of the plasticity rule for online learning from a perspective of objective function minimization due to performance requirements for robotic navigation applications. A variety of Hebbian plasticity rule formulations will lead to increasing the effective connectivity weights between co-active visual input feature neurons and compass neurons, but with different implicit objective functions that are being minimized in relation to network performance for orientation estimation. We propose a learning rule that is modulated by an additional variable (input current) and not purely homosynaptic to directly minimize orientation estimation error in the network which could be investigated experimentally.

There are several assumptions and simplifications in the presented results that could be investigated in future work. One simplification is that integration of a known linear velocity is utilized to perform path integration with an estimated orientation estimation. Future work could incorporate insect-inspired approaches for the path integration that assume an estimated orientation is already provided (*73*). Another simplification utilized in the network model is the processing of visual features by ring neurons with landmark specific tuning curves. We expect our findings to generalize across more visual feature encoding schemes such as more detailed models of the insect optic lobes, deep networks, or incorporation of processed dynamic vision sensor data. An additional area for model augmentation is to further constrain the ring attractor network with recently released synapse-level neural connectivity data (*59*).

For additional improvements to model performance, future enhancements could include further optimized weights between E-PG, P-EG, P-EN, and Pintr neurons. Further tuning the encoding of angular velocity including a mechanism for angular acceleration in P-EN inputs could improve the accuracy of angular integration and lead to an enhanced teaching signal for training visual feature inputs. Additionally, increasing the number or neurons or modeling neuronal compartments could further increase the number of stable states in the ring attractor. Direct evaluation of the proposed learning rule (Eq. 4), could be investigated in an additional series of simulation and robotic platform experiments. The performance of the network model could be further investigated further across longer trajectories and in multiple environments. A direct comparison of refined versions of the network model versus state-of-the-art benchmarking for lower power navigation approaches such as VIO will be critical to inform the ultimate viability of this approach for robotic translation.

We present a critical proof of principle for translation of insect-inspired approaches to robotics navigation to enable a future class of low SWaP algorithms to perform online learning and heading representation constrained in detail from biology. Given that all neurons are modeled as dynamic integrate-and-fire neurons, the model is amenable to incorporating event-driven low latency sensors such as dynamic vision sensors to enable updating estimates with visual features detected during high velocity movement. One of the key model features is the ability to utilize previously encountered visual features to update an estimate of orientation, loop closure for orientation estimation, which is not possible in low power navigation approaches utilizing VIO. While this is less than loop closure capabilities of a complete pose in full SLAM systems, it still is a promising functionality due to the ability for the model to be implemented as a parallelized distributed network on neuromorphic hardware. Additionally, compared to deep neural network approaches utilized in visual navigation where networks are trained offline due to computational resource and large data size requirements, the weights in the network model are learned online and are used over a single trajectory to increase estimation accuracy.

## Conclusion

In conclusion, we present a critical proof of concept for a low SWaP robotics navigation algorithm utilizing orientation estimation in a ring attractor network constrained using circuit details from *Drosophila* with online distributed learning amenable for neuromorphic implementation. By focusing on the objective function minimization necessary for a robotics implementation, we discovered a convenient formalism for common computational goals underlying both biological and artificial systems and identify testable predictions and areas of focus for future neuroscience experiments.

## Materials and Methods

All experiments are performed in a simulated or physical environment utilizing the same connectivity-constrained network model for performing orientation estimation.

### Network model input

A total of 81 ring neurons are simulated which are selectively tuned to the position of landmarks in the visual field in order to abstract the initial visual processing in the optic lobes. Specifically, sets of 27 ring neurons are selective to each of three landmarks with a Gaussian tuning curve with a standard deviation of 6.44° and a maximum response offset by 10° across a 270° field of view (Fig. 6A). The standard deviation of the Gaussian is selected such that adjacent neurons had overlapping turning curves starting at half maximum values. The current provided to each simulated E-PG neuron in each time step is determined by the visual ring neuron activation multiplied by ***W***^***r***→ ***c***^.

**Fig 6.**
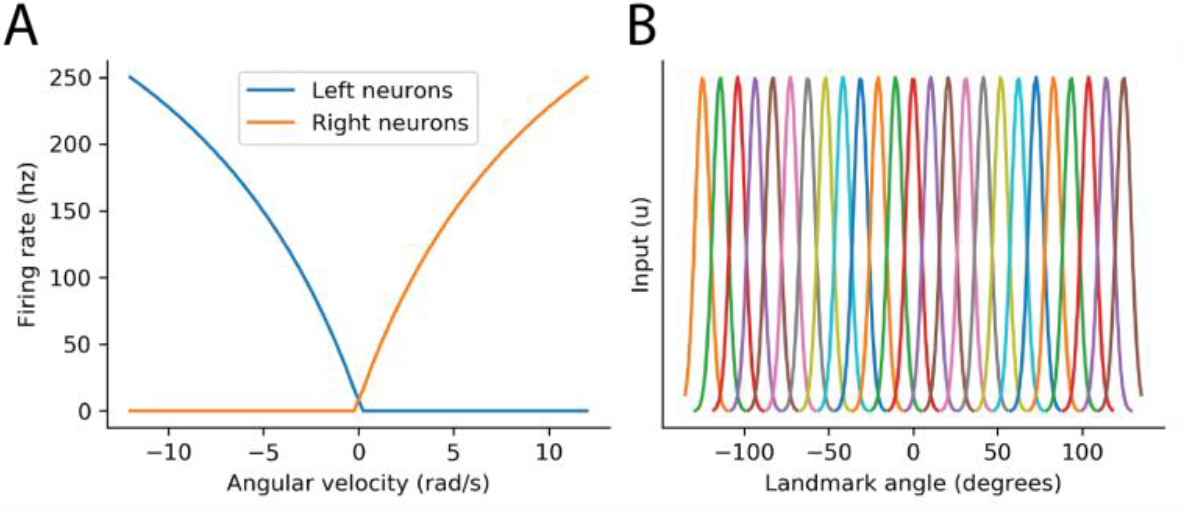
Model input encoding. (**A**) Receptive fields for each set of ring neurons over the field of view. (**B**) Angular velocity encoding in P-EN neurons.

The 16 P-EN neurons are split into two hemispheres (right and left), such that the 8 right P-EN neurons encode positive angular velocities, and the 8 left neurons negative velocities. The current provided to each neuron as input is calculated such that with no other input, each P-EN neuron has a steady state maximum firing rate of 250 Hz at 10 radians per second (Fig. 6B). In simulations, zero mean Gaussian noise is added to angular velocity with a standard deviation of 0.1 radians per second.

### Network model and configurations

Neural activation of each ring, E-PG, P-EG, P-EN, and Pintr neuron is modeled as leaky integrate and fire neurons utilizing the nengo software package (*75*) with a timestep of 1 millisecond. The external inputs driving the network activity are input currents from visual encoding to E-PG neurons and angular velocity encoding to P-EN neurons as described above. The connectivity pattern between E-PG, P-EG, P-EN, and Pintr neurons are based on previously reported biologically-constrained connectivity patterns (*67*). The network weights between neural subpopulations are 20 for all excitatory connections (E-PG→ P-EN, E-PG→ P-EG, E-PG→ Pintr, P-EN→ E-PG, P-EG→ E-PG), −15 for all inhibitory connections to excitatory neurons (Pintr→ P-EG, Pintr→ P-EN), and −20 for all Pintr→ Pintr connections. Stochasticity is introduced to the network with mean zero Gaussian noise with a standard deviation of 0.1 added to P-EG and Pintr neurons.

Three model configurations are used whose only difference is the weight of the ring neuron to E-PG connections. For the angular velocity only case, there is no visual input (the effective ***W***^***r***→ ***c***^ is **0**). For the offline optimal comparison, a supervised set of scaled ***W***^***r***→ ***c***^ weights is solved for using linear lasso regularized positive regression to minimize the objective function for each compass neuron 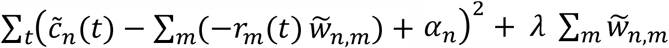, where 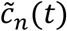 is a target set of compass neuron activation at each simulation timestep generated with the preferred angle of each compass neuron. In order to enforce negative weights for *w*_*n,m*_, the weights used in simulation are a scaled version of the 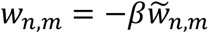, and ***β***=0.0025, to optimize estimation accuracy. For the online learning rule comparison, the weights between ring neurons and E-PG neurons are updated according to Eq. 1 with parameter values of 1.7e-6, 1.7e-4, and 0.29 for a, b, and *η*, respectively.

### Learning rule objective functions

In order to perform gradient descent on the objective function ***ϕ*** as defined in (Eq. 2) over time by modifying the synaptic weights, *w*_*nm*_, it follows from the chain rule that

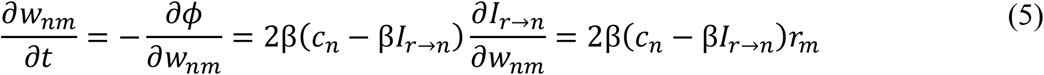

if we ignore the implicit dependence of *c*_*n*_ on *w*_*nm*_ assuming that the bump here is determined mostly by the recurrent weights. The discrete form of the gradient descent is

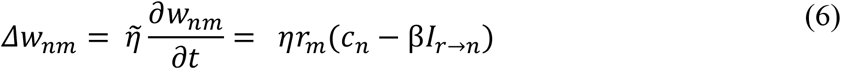

with a scalar learning rate parameter, 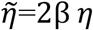. Similarly, to minimize the orientation error objective function, φ, as defined in (Eq. 3), it follows from the chain rule that

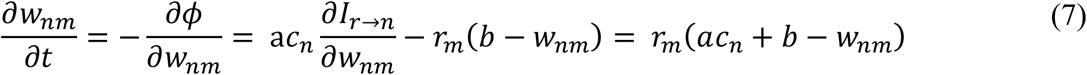

with a discrete form of

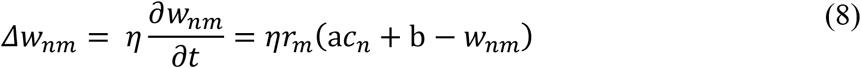

#### Hardware translation

The robotic platform utilized is a modified Robotis Turtlebot3 “Burger,” equipped with a Nvidia Jetson TX2, and a Ricoh Theta S 360° camera. The Turtlebot has an OpenCR embedded motor controller and sensor suite, as well as two Dynamixel XL430-W250 servos for wheel control.

The relative angle of colored landmarks in the arena are detected from 360° camera images utilizing color masks and blob detection prior to activation of visual ring neurons as described above with Gaussian receptive fields with respect to the relative angle of each landmark. All sensor data was recorded in ROS and used as input to the network model. We accelerated the input stream by a factor of ten to facilitate an evaluation of time courses corresponding to thos used in simulated environments.

The robot was run in an arena with an Optitrack system set up. IR-reflective markers were attached to the robot such that the position and orientation of the robot was tracked by the system. Red, green, blue, and yellow landmarks were made out of posterboard and placed in the environment with their own IR markers. Data from the Optitrack system was used for comparison.

## Author contributions

WGR, BR, KZ, EJ, and PR conceived and planned the research study and related experiments. BR, RT, KZ, and JJ performed network simulations and learning analysis. RT, MC, and DS performed robotic translation and related experiments. BR, RT, EJ, GH, KZ, and WG interpreted the results and wrote the manuscript. All authors provided critical feedback and helped shape the research, analysis and manuscript.

We would like to thank Michael Wolmetz and Joan Hoffmann for their insightful review and discussion in developing these experiments. We additionally would like to thank Kensei Suzuki, Lauren Diaz, Andres Perez-Doval, and Sonia Albert for their assistance with robotic translation, and the support of the JHU/APL CIRCUIT and ASPIRE programs.

## Disclaimer

This material is based upon work supported by (while GH was serving at) the National Science Foundation. Any opinion, findings, and conclusions or recommendations expressed in this material are those of the author(s) and do not necessarily reflect the views of the National Science Foundation.

## Data and materials availability

All data needed to support the conclusions in the paper are available via sources described in the paper or upon reasonable request to the authors.

## SUPPLEMENTARY MATERIALS

No supplementary materials are included.

## References and Notes

1. S. M. LaValle, Planning algorithms (Cambridge university press, 2006).

2. H. Durrant-Whyte, T. Bailey, Simultaneous localization and mapping: part I. IEEE Robot. Autom. Mag. 13, 99–110 (2006).

3. T. Bailey, H. Durrant-Whyte, Simultaneous localization and mapping (SLAM): Part II. IEEE Robot. Autom. Mag. 13, 108–117 (2006).

4. P. E. Hart, N. J. Nilsson, B. Raphael, A Formal Basis for the Heuristic Determination of Minimum Cost Paths. IEEE Trans. Syst. Sci. Cybern. 4, 100–107 (1968).

5. J. K. Uhlmann, Algorithms for multiple-target tracking. Am. Sci. 80, 128–141 (1992).

6. S. J. Julier, J. K. Uhlmann, in Signal processing, sensor fusion, and target recognition VI (1997), vol. 3068, pp. 182–193.

7. G. Loianno, C. Brunner, G. McGrath, V. Kumar, Estimation, control, and planning for aggressive flight with a small quadrotor with a single camera and IMU. IEEE Robot. Autom. Lett. 2, 404–411 (2016).

8. D. Scaramuzza, F. Fraundorfer, Visual odometry [tutorial]. IEEE Robot. Autom. Mag. 18, 80–92 (2011).

9. H. J. Kushner, Dynamical equations for optimal nonlinear filtering. J. Differ. Equ. 3, 179– 190 (1967).

10. P. M. Djuric, J. H. Kotecha, J. Zhang, Y. Huang, T. Ghirmai, M. F. Bugallo, J. Miguez, Particle filtering. IEEE Signal Process. Mag. 20, 19–38 (2003).

11. C. Hertzberg, R. Wagner, O. Birbach, T. Hammer, U. Frese, in 2011 IEEE International Conference on Robotics and Automation (2011), pp. 2644–2651.

12. M. Montemerlo, S. Thrun, D. Koller, B. Wegbreit, others, in IJCAI (2003), pp. 1151–1156.

13. R. Mur-Artal, J. M. M. Montiel, J. D. Tardos, ORB-SLAM: a versatile and accurate monocular SLAM system. IEEE Trans. Robot. 31, 1147–1163 (2015).

14. L. Zhao, S. Huang, G. Dissanayake, in 2013 IEEE/RSJ International Conference on Intelligent Robots and Systems (2013), pp. 24–30.

15. M. Abouzahir, A. Elouardi, R. Latif, S. Bouaziz, A. Tajer, Embedding SLAM algorithms: Has it come of age? Rob. Auton. Syst. 100, 14–26 (2018).

16. S. Aldegheri, N. Bombieri, D. D. Bloisi, A. Farinelli, in IEEE International Conference on Intelligent Robots and Systems (2019), pp. 5370–5375.

17. Q. Liu, S. Qin, B. Yu, J. Tang, S. Liu, π-BA: Bundle adjustment hardware accelerator based on distribution of 3d-point observations. IEEE Trans. Comput. 69, 1083–1095 (2020).

18. R. Li, S. Wang, D. Gu, Ongoing evolution of visual slam from geometry to deep learning: Challenges and opportunities. Cognit. Comput. 10, 875–889 (2018).

19. R. Li, S. Wang, Z. Long, D. Gu, in 2018 IEEE international conference on robotics and automation (ICRA) (2018), pp. 7286–7291.

20. A. Banino, C. Barry, B. Uria, C. Blundell, T. Lillicrap, P. Mirowski, A. Pritzel, M. J. Chadwick, T. Degris, J. Modayil, others, Vector-based navigation using grid-like representations in artificial agents. Nature. 557, 429–433 (2018).

21. S. K. Ramakrishnan, Z. Al-Halah, K. Grauman, Occupancy Anticipation for Efficient Exploration and Navigation. arXiv Prepr. arXiv:2008.09285 (2020).

22. R. S. Dimitrova, M. Gehrig, D. Brescianini, D. Scaramuzza, in 2020 IEEE International Conference on Robotics and Automation (ICRA) (IEEE, 2020; https://ieeexplore.ieee.org/document/9197530/), xpp. 4294–4300.

23. M. M. Khan, D. R. Lester, L. A. Plana, A. Rast, X. Jin, E. Painkras, S. B. Furber, in 2008 IEEE International Joint Conference on Neural Networks (IEEE World Congress on Computational Intelligence) (2008), pp. 2849–2856.

24. P. A. Merolla, J. V. Arthur, R. Alvarez-Icaza, A. S. Cassidy, J. Sawada, F. Akopyan, B. L. Jackson, N. Imam, C. Guo, Y. Nakamura, B. Brezzo, I. Vo, S. K. Esser, R. Appuswamy, B. Taba, A. Amir, M. D. Flickner, W. P. Risk, R. Manohar, D. S. Modha, A million spiking-neuron integrated circuit with a scalable communication network and interface. Science (80-.). 345, 668–673 (2014).

25. M. Davies, N. Srinivasa, T.-H. Lin, G. Chinya, Y. Cao, S. H. Choday, G. Dimou, P. Joshi, N. Imam, S. Jain, others, Loihi: A neuromorphic manycore processor with on-chip learning. IEEE Micro. 38, 82–99 (2018).

26. P. Blouw, X. Choo, E. Hunsberger, C. Eliasmith, in Proceedings of the 7th Annual Neuro-inspired Computational Elements Workshop (2019), pp. 1–8.

27. N. Imam, T. A. Cleland, Rapid online learning and robust recall in a neuromorphic olfactory circuit. Nat. Mach. Intell. 2, 181–191 (2020).

28. J. S. Taube, The Head Direction Signal: Origins and Sensory-Motor Integration. Annu. Rev. Neurosci. 30, 181–207 (2007).

29. E. I. Moser, E. Kropff, M. B. Moser, Place cells, grid cells, and the brain’s spatial representation system. Annu. Rev. Neurosci. 31, 69–89 (2008).

30. J. O’Keefe, Place units in the hippocampus of the freely moving rat. Exp. Neurol. 51, 78– 109 (1976).

31. S. Poulter, T. Hartley, C. Lever, The Neurobiology of Mammalian Navigation. Curr. Biol. 28 (2018), pp. R1023–R1042.

32. J. D. Monaco, R. M. De Guzman, H. T. Blair, K. Zhang, Spatial synchronization codes from coupled rate-phase neurons. PLoS Comput. Biol. 15 (2019), doi:10.1371/journal.pcbi.1006741.

33. M. J. Milford, G. F. Wyeth, D. Prasser, in IEEE International Conference on Robotics and Automation, 2004. Proceedings. ICRA’04. 2004 (2004), vol. 1, pp. 403–408.

34. J. Steckel, H. Peremans, BatSLAM: Simultaneous localization and mapping using biomimetic sonar. PLoS One. 8, e54076 (2013).

35. A. Arleo, W. Gerstner, in From Animals to Animats 6 (The MIT Press, 2000).

36. R. Kreiser, A. Renner, Y. Sandamirskaya, P. Pienroj, Pose Estimation and Map Formation with Spiking Neural Networks: Towards Neuromorphic SLAM. IEEE Int. Conf. Intell. Robot. Syst., 2159–2166 (2018).

37. G. Tang, K. P. Michmizos, in ACM International Conference Proceeding Series (2020).

38. J. D. Monaco, G. M. Hwang, K. M. Schultz, K. Zhang, Cognitive swarming in complex environments with attractor dynamics and oscillatory computing. Biol. Cybern. 114, 269– 284 (2020).

39. G. M. Muir, J. E. Brown, J. P. Carey, T. P. Hirvonen, C. C. Della Santina, L. B. Minor, J. S. Taube, Disruption of the head direction cell signal after occlusion of the semicircular canals in the freely moving chinchilla. J. Neurosci. 29, 14521–14533 (2009).

40. R. G. Robertson, E. T. Rolls, P. Georges-François, S. Panzeri, Head direction cells in the primate pre-subiculum. Hippocampus. 9, 206–219 (1999).

41. A. Finkelstein, D. Derdikman, A. Rubin, J. N. Foerster, L. Las, N. Ulanovsky, Three-dimensional head-direction coding in the bat brain. Nature. 517, 159–164 (2015).

42. J. D. Seelig, V. Jayaraman, Neural dynamics for landmark orientation and angular path integration. Nature. 521, 186–191 (2015).

43. W. E. Skaggs, J. J. Knierim, H. S. Kudrimoti, B. L. McNaughton, A model of the neural basis of the rat’s sense of direction. Adv. Neural Inf. Process. Syst. 7, 173–180 (1995).

44. A. D. Redish, A. N. Elga, D. S. Touretzky, A coupled attractor model of the rodent head direction system. Netw. Comput. Neural Syst. 7, 671–685 (1996).

45. K. Zhang, Representation of spatial orientation by the intrinsic dynamics of the head-direction cell ensemble: A theory. J. Neurosci. 16, 2112–2126 (1996).

46. X. Xie, R. H. R. Hahnloser, H. S. Seung, Double-ring network model of the head-direction system. Phys. Rev. E − Stat. Physics, Plasmas, Fluids, Relat. Interdiscip. Top. 66, 9 (2002).

47. S. M. Stringer, T. P. Trappenberg, E. T. Rolls, I. E. T. d. Araujo, Self-organizing continuous attractor networks and path integration: one-dimensional models of head direction cells. Netw. Comput. Neural Syst. 13, 217–242 (2002).

48. C. Boucheny, N. Brunel, A. Arleo, A continuous attractor network model without recurrent excitation: Maintenance and integration in the head direction cell system. J. Comput. Neurosci. 18, 205–227 (2005).

49. P. Song, X. J. Wang, Angular path integration by moving “hill of activity”: A spiking neuron model without recurrent excitation of the head-direction system. J. Neurosci. 25, 1002–1014 (2005).

50. P. E. Sharp, H. T. Blair, J. Cho, The anatomical and computational basis of the rat head-direction cell signal. Trends Neurosci. 24, 289–294 (2001).

51. K. E. Cullen, J. S. Taube, Our sense of direction: Progress, controversies and challenges. Nat. Neurosci. 20, 1465–1473 (2017).

52. T. M. Massoud, T. K. Horiuchi, A Neuromorphic VLSI Head Direction Cell System. IEEE Trans. Circuits Syst. I Regul. Pap. 58, 150–163 (2011).

53. R. Kreiser, M. Cartiglia, J. N. P. Martel, J. Conradt, Y. Sandamirskaya, A Neuromorphic Approach to Path Integration: A Head-Direction Spiking Neural Network with Vision-driven Reset. Proc. - IEEE Int. Symp. Circuits Syst. 2018–May (2018).

54. T. Degris, C. Boucheny, A. Arleo, A. Lip, C. Scott, in From Animals to Animats 8 (The MIT Press, 2004).

55. T. Kyriacou, Using an evolutionary algorithm to determine the parameters of a biologically inspired model of head direction cells. J. Comput. Neurosci. 32, 281–295 (2012).

56. G. Tejera, M. Llofriu, A. Barrera, A. Weitzenfeld, Bio-Inspired Robotics: A Spatial Cognition Model integrating Place Cells, Grid Cells and Head Direction Cells. J. Intell. Robot. Syst. Theory Appl. 91, 85–99 (2018).

57. X. Zhou, C. Weber, S. Wermter, in Lecture Notes in Computer Science (including subseries Lecture Notes in Artificial Intelligence and Lecture Notes in Bioinformatics) (2017), vol. 10613 LNCS, pp. 137–145.

58. A. Bicanski, N. Burgess, Environmental anchoring of head direction in a computational model of retrosplenial cortex. J. Neurosci. 36, 11601–11618 (2016).

59. L. K. Scheffer, C. S. Xu, M. Januszewski, Z. Lu, S. Takemura, K. J. Hayworth, G. B. Huang, K. Shinomiya, J. Maitlin-Shepard, S. Berg, J. Clements, P. M. Hubbard, W. T. Katz, L. Umayam, T. Zhao, D. Ackerman, T. Blakely, J. Bogovic, T. Dolafi, D. Kainmueller, T. Kawase, K. A. Khairy, L. Leavitt, P. H. Li, L. Lindsey, N. Neubarth, D. J. Olbris, H. Otsuna, E. T. Trautman, M. Ito, A. S. Bates, J. Goldammer, T. Wolff, R. Svirskas, P. Schlegel, E. Neace, C. J. Knecht, C. X. Alvarado, D. A. Bailey, S. Ballinger, J. Borycz, B. S. Canino, N. Cheatham, M. Cook, M. Dreher, O. Duclos, B. Eubanks, K. Fairbanks, S. Finley, N. Forknall, A. Francis, G. P. Hopkins, E. M. Joyce, S. Kim, N. A. Kirk, J. Kovalyak, S. A. Lauchie, A. Lohff, C. Maldonado, E. A. Manley, S. McLin, C. Mooney, M. Ndama, O. Ogundeyi, N. Okeoma, C. Ordish, N. Padilla, C. M. Patrick, T. Paterson, E. E. Phillips, E. M. Phillips, N. Rampally, C. Ribeiro, M. K. Robertson, J. T. Rymer, S. M. Ryan, M. Sammons, A. K. Scott, A. L. Scott, A. Shinomiya, C. Smith, K. Smith, N. L. Smith, M. A. Sobeski, A. Suleiman, J. Swift, S. Takemura, I. Talebi, D. Tarnogorska, E. Tenshaw, T. Tokhi, J. J. Walsh, T. Yang, J. A. Horne, F. Li, R. Parekh, P. K. Rivlin, V. Jayaraman, M. Costa, G. S. Jefferis, K. Ito, S. Saalfeld, R. George, I. A. Meinertzhagen, G. M. Rubin, H. F. Hess, V. Jain, S. M. Plaza, A connectome and analysis of the adult Drosophila central brain. Elife. 9 (2020), doi:10.7554/eLife.57443.

60. F. Li, J. Lindsey, E. C. Marin, N. Otto, M. Dreher, G. Dempsey, I. Stark, A. Shakeel Bates, M. William Pleijzier, P. Schlegel, S. Takemura, T. Yang, A. Francis, A. Braun, R. Parekh, M. Costa, L. Scheffer, Y. Aso, G. S. X E Jefferis, L. Abbott, S. Waddell, G. M. Rubin, bioRxiv, doi:10.1101/2020.08.29.273276.

61. S. S. Kim, H. Rouault, S. Druckmann, V. Jayaraman, Ring attractor dynamics in the Drosophila central brain. Science (80-.). 356, 849–853 (2017).

62. D. Turner-Evans, S. Wegener, H. Rouault, R. Franconville, T. Wolff, J. D. Seelig, S. Druckmann, V. Jayaraman, Angular velocity integration in a fly heading circuit. Elife. 6, 1–39 (2017).

63. J. Green, A. Adachi, K. K. Shah, J. D. Hirokawa, P. S. Magani, G. Maimon, A neural circuit architecture for angular integration in Drosophila. Nat. Publ. Gr. 546, 101–106 (2017).

64. J. D. Seelig, V. Jayaraman, Feature detection and orientation tuning in the Drosophila central complex. Nature. 503, 262–266 (2013).

65. Y. E. Fisher, J. Lu, I. D’Alessandro, R. I. Wilson, Sensorimotor experience remaps visual input to a heading-direction network. Nature. 576 (2019).

66. T. Wolff, N. A. Iyer, G. M. Rubin, Neuroarchitecture and neuroanatomy of the Drosophila central complex: A GAL4-based dissection of protocerebral bridge neurons and circuits. J. Comp. Neurol. 523, 997–1037 (2015).

67. K. S. Kakaria, B. L. de Bivort, Ring Attractor Dynamics Emerge from a Spiking Model of the Entire Protocerebral Bridge. Front. Behav. Neurosci. 11, 1–13 (2017).

68. T. S. Su, W. J. Lee, Y. C. Huang, C. Te Wang, C. C. Lo, Coupled symmetric and asymmetric circuits underlying spatial orientation in fruit flies. Nat. Commun. 8 (2017), doi:10.1038/s41467-017-00191-6.

69. I. Pisokas, S. Heinze, B. Webb, The head direction circuit of two insect species. Elife. 9, 1– 49 (2020).

70. A. J. Cope, C. Sabo, E. Vasilaki, A. B. Barron, J. A. R. Marshall, A computational model of the integration of landmarks and motion in the insect central complex. PLoS One. 12, 1– 19 (2017).

71. S. S. Kim, A. M. Hermundstad, S. Romani, L. F. Abbott, V. Jayaraman, Generation of stable heading representations in diverse visual scenes. Nature. 576 (2019).

72. Z. Zhang, X. Li, J. Guo, Y. Li, A. Guo, Two clusters of GABAergic ellipsoid body neurons modulate olfactory labile memory in Drosophila. Ann. Intern. Med. 158, 5175–5181 (2013).

73. T. Stone, B. Webb, A. Adden, N. Ben Weddig, A. Honkanen, R. Templin, W. Wcislo, L. Scimeca, E. Warrant, S. Heinze, An Anatomically Constrained Model for Path Integration in the Bee Brain. Curr. Biol. 27, 3069–3085.e11 (2017).

74. G. Klančar, A. Zdešar, S. Blažič, I. Škrjanc, Wheeled Mobile Robotics: From Fundamentals Towards Autonomous Systems (2017).

75. T. Bekolay, J. Bergstra, E. Hunsberger, T. DeWolf, T. C. Stewart, D. Rasmussen, X. Choo, R. Voelker, C. Eliasmith, Nengo: A Python tool for building large-scale functional brain models. Front. Neuroinform. 7, 1–13 (2014).

